# Protein Binding Leads to Reduced Stability and Solvated Disorder in the Polystyrene Nanoparticle Corona

**DOI:** 10.1101/2023.07.06.548033

**Authors:** Radha P. Somarathne, Dhanush L. Amarasekara, Chathuri S. Kariyawasam, Harley A. Robertson, Railey Mayatt, Nicholas C. Fitzkee

**Affiliations:** Department of Chemistry, Mississippi State University, Mississippi State, MS 39762 USA

**Keywords:** nanoparticle, protein, corona, structure, interactions

## Abstract

Understanding the conformation of proteins in the nanoparticle corona has important implications in how organisms respond to nanoparticle-based drugs. These proteins coat the nanoparticle surface, and their properties will influence the nanoparticle’s interaction with cell targets and the immune system. While some coronas are thought to be disordered, two key unanswered questions are the degree of disorder and solvent accessibility. Here, using a comprehensive thermodynamic approach, along with supporting spectroscopic experiments, we develop a model for protein corona disorder in polystyrene nanoparticles of varying size. For two different proteins, we find that binding affinity decreases as nanoparticle size increases. The stoichiometry of binding, along with changes in the hydrodynamic size, support a highly solvated, disordered protein corona anchored at a small number of enthalpically-driven attachment sites. The scaling of the stoichiometry vs. nanoparticle size is consistent disordered polymer dimensions. Moreover, we find that proteins are destabilized less severely in the presence of larger nanoparticles, and this is supported by measurements of hydrophobic exposure, which becomes less pronounced at lower curvatures. Our observations hold for flat polystyrene surfaces, which, when controlled for total surface area, have the lowest hydrophobic exposure of all systems. Our model provides an explanation for previous observations of increased amyloid fibrillation rates in the presence of larger nanoparticles, and it may rationalize how cell receptors can recognize protein disorder in therapeutic nanoparticles.

**TOC Image:** 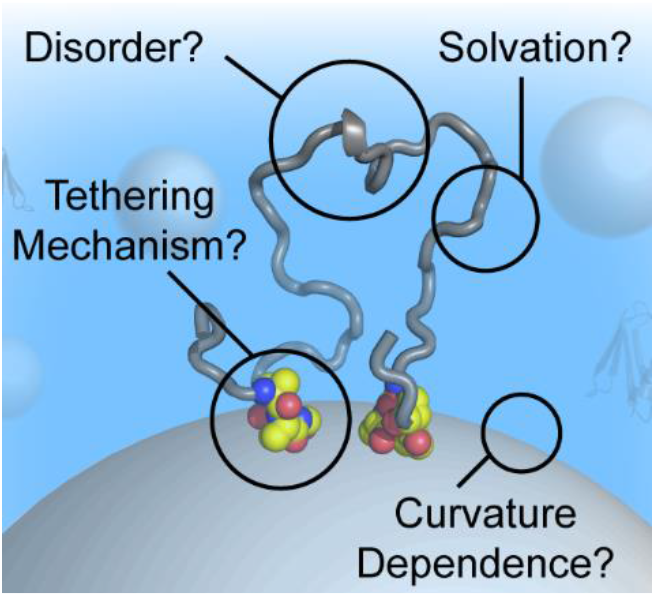

## Introduction

Nanotechnology is an invaluable tool with versatile applications in medicine, sensing, and biotechnology.^1–3^ Tailoring nanoparticle interactions with their environment is a critical aspect of these applications. In biological fluids, protein adsorption to surfaces of nanomaterials is a widely known phenomenon and can pose a significant challenge. The layer of adsorbed proteins is known as the “protein corona,” and it remains a research topic of high importance.^4,5^ The corona is often characterized as either “hard” or “soft,” depending on the rate at which nanoparticle-associated proteins can exchange with the solution.^5–7^ More dynamic exchange is characteristic of a soft corona, whereas long-lived adsorption (> min) is characteristic of a “hard” corona. The importance of the corona cannot be understated. A recent literature review implicated the corona as the reason why, for an injected dose of a typical nanoparticle-based therapeutic, only 1.5-2.2% of nanoparticles reach their intended targets.^8^ The corona has been shown to influence nanoparticle uptake,^1,9^ in vivo pharmacokinetics, and biodistribution,^10,11^ and it represents what the immune system and cell targets “see” in their biological context. Presumably, the structure of adsorbed proteins influences how nanoparticles are trafficked, e.g., through interactions with complement proteins.^12^ Therefore, understanding the structure and organization of proteins in the corona is crucial.

Currently, little is understood about the structure of these corona proteins. Mass spectrometry-based proteomics studies have revealed a great deal about the identity of adsorbed proteins,^13,14^ and advances in chromatography and spectroscopy have enabled measurements of kinetic and thermodynamic binding parameters.^15–18^ However, little is known about the structure of proteins once they adsorb to the surface. A recent study by Zhang *et al.* used antibodies to probe the accessible epitopes on the surface of gold nanoparticles (AuNPs).^19^ Using a matrix of pairwise interactions, they found that densely packed proteins in the corona can alter protein structure and epitope accessibility. Our recent work, also on AuNPs, identified that protein orientation and function could be successfully predicted in many cases using a simple algorithm that quantifies each amino acid’s interaction with a gold surface.^20^ However, since adsorption exhibits aspects of kinetic control,^21^ there is significant heterogeneity in the corona. For example, cryo-electron microscopy reveals a fairly inhomogeneous protein density around the AuNP surface with no clear patterns or organization.^22^ Thus, while a particular orientation may be preferred, the nanoparticle corona is not uniform, and proteins sample numerous conformations on the surface. This is likely especially true for proteins in complex mixtures such as blood serum. Moreover, all of these studies were performed on AuNPs, which are known to have a relatively hard, non-exchangeable corona.^4^ Such a corona simplifies the characterization of long-lived adsorbed proteins; proteins in the soft corona are much harder to study because of their dynamic exchange.

Nanoparticle curvature has proven to be a useful tool in understanding adsorbed protein properties. Altering nanoparticle size and curvature changes the geometry of the surface without altering its chemical composition, and this can be useful when investigating protein stability and structure.^7,23,24^ For example, early work on gold nanospheres and nanorods demonstrated that both colloidal stability and adsorbed enzyme function could be altered depending on the size and shape of the AuNPs.^25,26^ On silica nanospheres, it was shown that adsorbed protein structure varied depending on the packing density and whether the protein was globular; albumin tended to lose its structure on larger particles, whereas fibrinogen was stabilized by side-on interactions present on larger particles.^27^ Mandal and Kraatz observed that, for self-assembled monolayers of small helical peptides, highly curved surfaces on 5 nm AuNPs were most destabilizing, but peptide structure was unperturbed for 20 nm AuNPs and even flat gold surfaces. This demonstrates that a range of stabilities are expected depending on the curvature, but the structural properties of larger proteins remain unclear. Often, studies involving nanoparticle curvature are not controlled for total surface area, which may confound interpretations of protein structure, especially if the binding affinity changes and surface saturation cannot be ensured. Clearly, however, side-on interactions play a role in corona stabilization,^27^ and there is much to be learned from studies covering a range of surface curvatures.

Overall, most investigations of adsorbed protein structural properties have been limited to broad statements about whether function is retained, whether the protein is globular, and the average orientation.^4,20,28–31^ If proteins are disordered, however, it is not always clear whether the disordered proteins are densely packed, extended, or well-solvated. Since a significant number of protein corona studies implicate protein disorder,^32,33^ it is important to understand the degree to which disorder is present and disordered protein accessibility. These factors will strongly influence interactions with the immune system and other biological processes that determine nanoparticle fate.

In this work, we use rigorous thermodynamic approaches to understand the structural properties of proteins adsorbed to polystyrene nanoparticles (PSNPs). These spherical particles exhibit a reversibly bound, soft corona and are thought to destabilize adsorbed proteins,^34–36^ but it is not clear whether the adsorbed state behaves like a traditional unfolded protein or something else. By using carefully controlled calorimetric titrations along with protein stability measurements for two different proteins, we determine the degree to which proteins are destabilized on PSNPs of varying curvature. The stoichiometry, combined with observed changes in hydrodynamic diameter, suggest that adsorbed proteins are highly extended on the surface with significant solvent exposure. In addition, the trend in surface coverage is consistent with what would be expected for a well-hydrated polymer. These observations, combined with circular dichroism (CD) and measurements of hydrophobic exposure, indicate that the nanoparticle attachment sites are limited, and that a majority of an adsorbed protein on smaller PSNPs (50 nm) is accessible to solvent or other proteins. The trends in stability can be extrapolated to flat polystyrene surfaces, where proteins appear to have minimal hydrophobic exposure. Using our data, we propose a model for how immune system components can recognize disorder in nanoparticle coronas, and we explain why nanoparticles are occasionally observed to induce fibrils and aggregates.

### Experimental Section

#### Protein expression and purification

Expression and purification of R2ab and GB3 were performed as described previously.^36,37^ Purified proteins were lyophilized in water (R2ab) or flash frozen in liquid nitrogen (GB3) and stored at -80 °C until they were ready to use.

#### Nanoparticle preparation and characterization

Non-functionalized polystyrene nanoparticles with varying diameters of 50 nm, 100 nm, and 200 nm were purchased from PolySciences (#08691-10, #00876-15, and #07304-15, respectively). Manufacturer-listed nanoparticle diameters were confirmed by dynamic light scattering (DLS). The manufacturer-supplied concentration was used, assuming no losses during dialysis or dilution; initial gravimetric analysis confirmed a concentration within 5% of the manufacturer’s stated value.

DLS measurements were performed in a working buffer (20 mM NaH_2_PO_4_ pH 6.5, 50 mM NaCl) to determine the hydrodynamic diameters of particles together with the polydispersity index (PDI). Additionally, the particles’ zeta potential was determined to investigate colloidal stability. These measurements were obtained using an Anton Paar Litesizer 500 DLS instrument at 25 °C, and the data was processed using the Kalliope software. Before mixing with proteins, all nanoparticles were dialyzed extensively in water to remove any traces of SDS. Absence of SDS was confirmed using NMR and contact angle measurements.^38^ A typical dialysis involved three exchanges into 1L of milli-Q water, allowing exchange to occur for at least four hours each time. Unless otherwise stated, PSNP concentrations were adjusted to maintain a constant surface area of 50 cm^2^. The protein concentrations and nanoparticle concentrations are listed in the Supporting Information (**Table S1**).

To confirm the PSNP size, 5 μL of 2 nM PSNP solution was deposited on Formvar-coated copper grids (Electron Microscopy Sciences, #FCF300-CU). The excess liquid was wicked away, and the remaining thin film on the grid was allowed to dry for 24 h at room temperature. The prepared grids were imaged using a JEOL 2100 TEM with an accelerating voltage of 200 kV to obtain clear images of well-dispersed nanoparticles (Supporting Information, **Fig. S1**). Average size was determined using the ImageJ software suite.

#### Isothermal titration calorimetry (ITC)

ITC measurements were performed on a VP-ITC (GE Healthcare). All protein and nanoparticle solutions were dialyzed in 20 mM NaH_2_PO_4_ and 50 mM NaCl buffer at pH 6.5. The proteins were titrated into nanoparticle dispersions at various concentrations to achieve titration curves with a clear sigmoidal transition. A typical titration consisted of 28 injections of 10 μL of protein solution being injected, with an equilibration time of 5 minutes between injections to ensure the recovery of the baseline before the next injection. Typical concentrations for 50 nm PSNPs were 25 μM protein in the syringe and 150 nM PSNPs in the ITC cell. During each experiment, the cell was continuously stirred at 480 rpm and the cell temperature was maintained at 25 °C. Representative titrations for all combinations of proteins and PSNPs are provided (**Fig. 1** and Supporting Information, **Fig. S2**). Control experiments were performed to determine the heat of the dilution of particles and proteins into the buffer, which were small (Supporting Information, **Fig. S3**). Titration of water from the syringe into the cell containing water was also performed regularly to ensure that the ITC system was operating optimally. The integrated heats for the titration of protein into particles were calculated and corrected with the corresponding heats of dilution in the buffer. Baseline corrections for the raw thermograms were performed using NITPIC to avoid introducing bias.^39^ The data was analyzed using Origin software and CHASM^40^ to obtain the binding stoichiometry of the complexes. The single set of identical sites model or the two sets of identical sites model was used depending on the profile observed. All ITC experiments were performed in triplicate (or more) using independently prepared replicate samples. Uncertainties in *N*, *K*, Δ_*bind*_*H*, Δ_*bind*_*G*^*o*^, and −*T*Δ_*bind*_*S* are reported as the standard error of the mean from these replicates.

**Figure 1.**
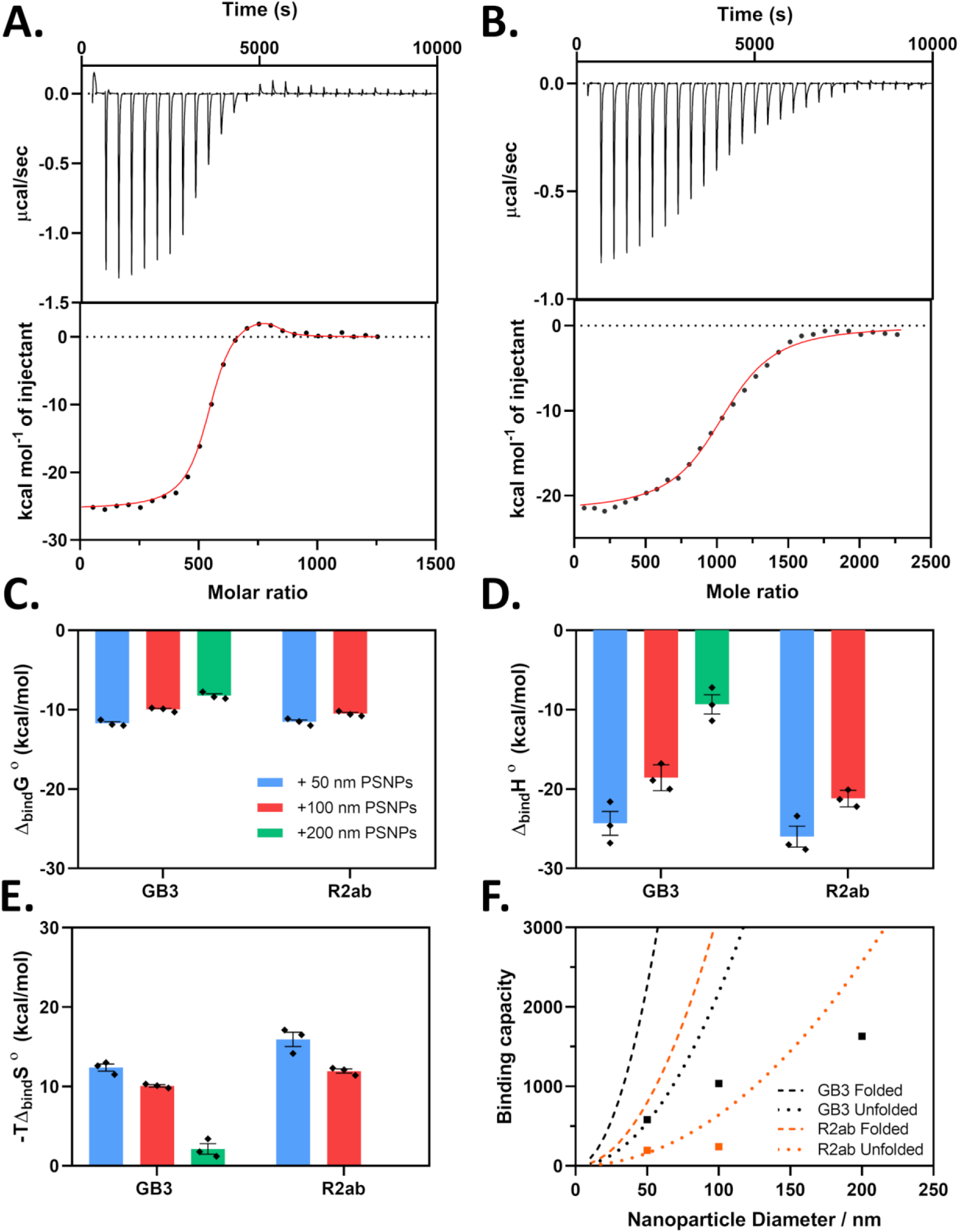
Representative ITC data for GB3 bound to 50 nm (A) and 100 nm PSNPs (B) is shown. Binding interactions in the nanoparticle corona weaken as the nanoparticle size increases. ITC parameters are shown for GB3 and R2ab in the presence of 50 nm (blue), 100 nm (red) and 200 nm (green) PSNPs. Free energies (C), and binding enthalpies (D) decrease as PSNP size increases. The entropy change also decreases, making the reaction more favorable on larger PSNPs (E). A comparison of experimental (solid squares) and expected N values for nanoparticles as put forth by Kohn et. al.is shown in (D). The error bars for each data point represent the standard error of the mean.

#### ANS fluorescence

Fluorescence measurements with 8-anilino-1-naphthalenesulfonic acid (ANS, Invitrogen #A47) were performed using a Spectramax 4 plate reader (Molecular Devices) at 25 °C. The final ANS concentration in each sample was 10 μM unless stated otherwise. The final GB3 and R2ab concentration were 20 and 7 μM, respectively. For mixtures of PSNPs, concentrations were set to ensure saturation of the surface and maintain a constant total nanoparticle surface area. The PSNP concentrations used were 8.8, 2.2 and 0.6 nM for 50, 100, and 200 nm PSNPs, respectively. The fluorescence intensities were measured at an excitation wavelength of 380 nm and the fluorescence signal was scanned from 400 nm to 650 nm. The blank samples included only PSNPs, buffer, and ANS. Protein-containing samples were incubated at room temperature for one hour in the dark, and the blank spectrum was subtracted from the spectrum of protein, PSNPs, buffer, and ANS. A blank containing PSNPs and ANS was necessary because ANS exhibits fluorescence in the presence of PSNPs but in the absence of proteins (Supporting Information, **Fig. S4**). Once protein binding occurred, however, the fluorescence signal was stable up to 12 hours after mixing (Supporting Information, **Fig. S5**).

Experiments involving flat polystyrene plates used a lower volume and a lower concentration of PSNPs. For a 34.8 mm well in a 6-well polystyrene plate (Corning #3516), the approximate surface area of the bottom surface is 9.51 cm^2^; final PSNP concentrations were determined to match this surface area (Supporting Information, **Table S2**). This plate was filled to a final volume of 0.5 mL to minimize contact with the walls of the well, and the solutions were agitated to ensure coverage over the entire bottom surface. The final ANS concentration in each sample was 10 μM, and the final GB3 and R2ab concentration were 1 and 0.3 μM, respectively. The samples were excited at 380 nm and the emission was observed at 480 nm. Each well was prepared independently in triplicate, with the appropriate blanks as described in the text. When applicable, the individual data points are reported in each figure, and the uncertainties are calculated as the standard error of the mean for these independent samples.

#### Intrinsic Tryptophan Fluorescence

Tryptophan fluorescence emission spectra for proteins in the presence of PSNPs of all sizes were monitored using a Horiba Fluoromax 4 fluorescence spectrophotometer equipped with a Hamilton Microlab 600 Titrator and Quantum-Northwest Peltier temperature control. Guanidinium chloride (GdmCl, MP Biomedicals #820539) solutions were prepared with a typical a stock concentration of 6.8 M. The actual GdmCl concentration for each sample was determined using refractometry.^41^ Unfolding experiments were conducted by monitoring the intrinsic tryptophan fluorescence as a function of increasing GdmCl concentration. To minimize the contribution from Tyr residues, samples were excited at 295 nm. The slit width for these studies 3 nm. The constant volume method was used to keep the overall cuvette volume constant, and the denaturant titrations were performed while maintaining an equilibration time of 6 minutes in between injections. The cuvette was subjected to constant stirring throughout the experiment, to maintain sample homogeneity. The final spectra are corrected for protein dilution, and each measurement was performed in triplicate from three or more independently-prepared samples. Titrations were fit to the linear extrapolation model using a Python-based fitting routine.^42^ All thermodynamic parameters are reported as the average and standard error of the mean.

#### Far – UV Circular Dichroism

The circular dichroism (CD) measurements were carried out using a Jasco 1500 CD spectrometer at 25 °C. The measurements were performed at a path length of 0.2 mm using a quartz cuvette. To determine how the secondary structure of both proteins changed in the presence of nanoparticles, all the protein solutions were made to contain the same concentration of proteins and nanoparticles used in the ANS fluorescence experiments in polystyrene plates. The buffer for all solutions was 10 mM sodium phosphate (pH 7.0). Between each sample measurement, the cuvette was carefully cleaned. Far-UV spectra were collected between 180 to 260 nm, with the scan rate set at 5 nm /min at a bandwidth of 1 nm using 16 s as the integration time. The data was analyzed and transformed into molar ellipticity using Jasco’s Spectra Manager software suite.

## Results

### A Sizeable Protein Corona Forms on Highly Curved Polystyrene Nanoparticle Surfaces

We selected two proteins for an investigation of energetics in the polystyrene nanoparticle protein corona: The first, GB3, is a small model protein that has been studied extensively on metallic nanoparticles.^4,15,43–45^ The second, R2ab, is implicated in the initial bacterial attachment during biofilm formation,^46,47^ and it is known to interact strongly with polystyrene surfaces.^36^ In the absence of protein, all nanoparticles were well dispersed and homogenous in size. The observed hydrodynamic diameters of bare nanoparticles agreed with the manufacturer’s specifications, and lot-to-lot values were consistent with the reported uncertainty (**Table 1**). TEM was also used to characterize bare PSNPs (Supporting Information, **Fig. S1**). The zeta potential (ζ) of the nanoparticles, which reflects the electrophoretic mobility, was negative for all particles observed (Supporting Information, **Table S3**). This negative surface charge originates from sulfonate groups present on the surface of the nanoparticles, and these groups are introduced during the polymerization process.^48^

**Table 1.**
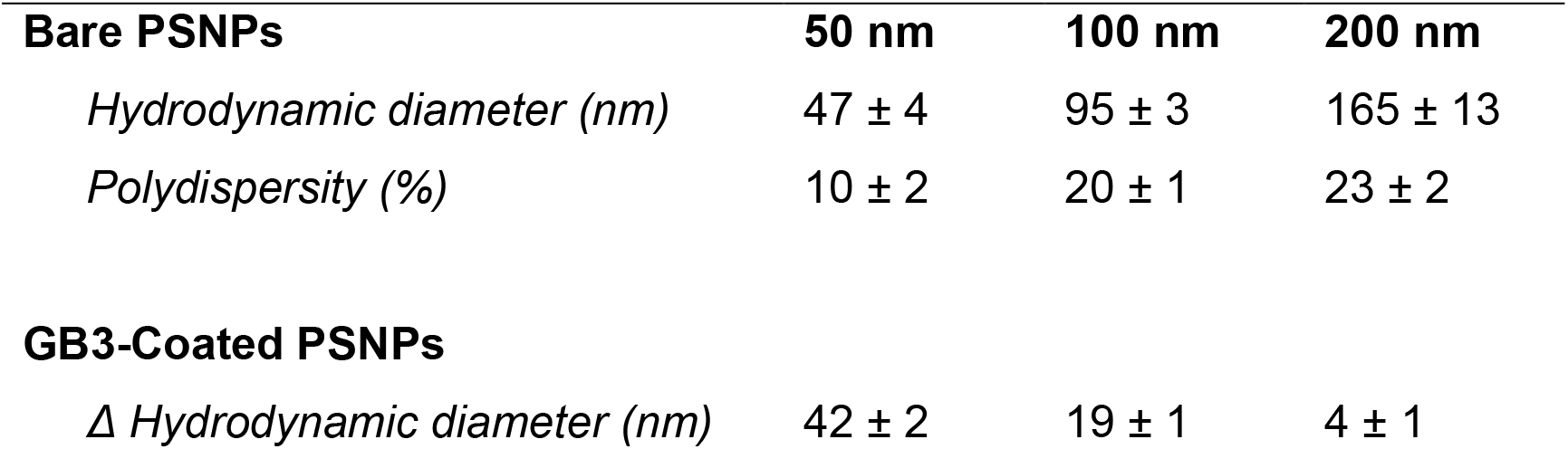
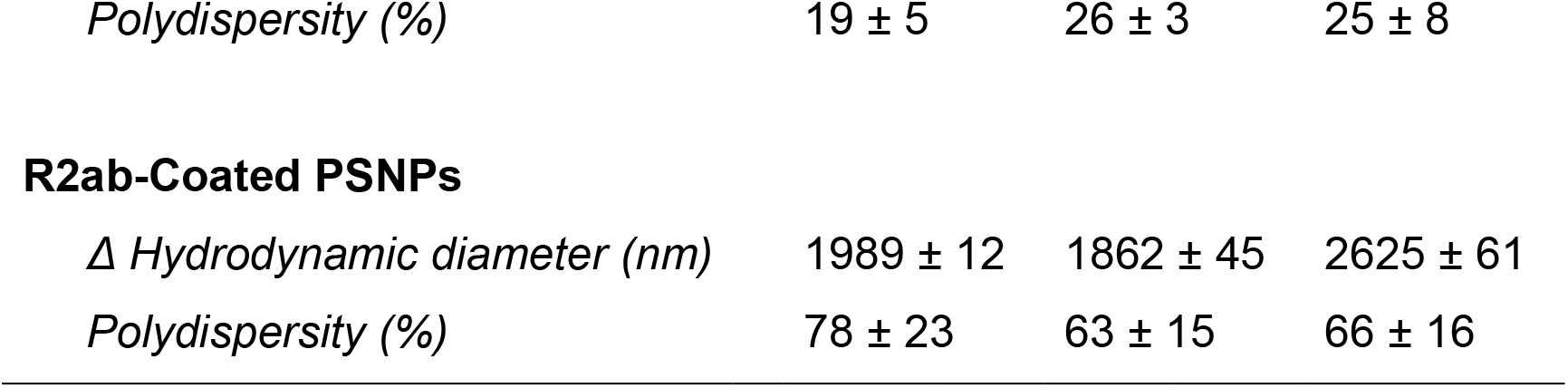
Hydrodynamic diameters of PSNPs and the observed increase in size upon interaction with GB3 and R2ab.

| **Bare PSNPs** | **50 nm** | **100 nm** | **200 nm** |
| --- | --- | --- | --- |
| *Hydrodynamic diameter (nm)* | 47 ± 4 | 95 ± 3 | 165 ± 13 |
| *Polydispersity (%)* | 10 ± 2 | 20 ± 1 | 23 ± 2 |
| **GB3-Coated PSNPs** |  |  |  |
| *$\Delta$ Hydrodynamic diameter (nm)* | 42 ± 2 | 19 ± 1 | 4 ± 1 |
| *Polydispersity (%)* | $19 \pm 5$ | $26 \pm 3$ | $25 \pm 8$ |
| **R2ab-Coated PSNPs** |  |  |  |
| $\Delta$ *Hydrodynamic diameter (nm)* | $1989 \pm 12$ | $1862 \pm 45$ | $2625 \pm 61$ |
| *Polydispersity (%)* | $78 \pm 23$ | $63 \pm 15$ | $66 \pm 16$ |

In the presence of GB3, an increase is in the hydrodynamic diameter is observed for all nanoparticles. The polydispersity remains relatively low (< 25%), indicating that the GB3 corona on PSNPs does not significantly affect the nanoparticles’ colloidal stability. However, as the nanoparticle size is increased, the increase in hydrodynamic diameter from the protein corona diminishes (**Table 1**). For 50 nm particles, the GB3 corona nearly doubles the apparent hydrodynamic diameter, from 47 ± 4 to 89 ± 5 (Δ*d*_*H*_ = 42 ± 6). However, for 200 nm PSNPs, the observed Δ*d*_*H*_ is substantially smaller, only 4 ± 1 nm. The observed increase for 100 nm PSNPs is intermediate (19 ± 1 nm). For all of these cases, we dialyzed the nanoparticles so that the solution conditions were consistent, and we maintained a constant ratio between protein concentration to total PSNP surface area. This finding suggests that the energetics induced by nanoparticle curvature differ significantly between 50 and 200 nm nanoparticles. GB3 has a standard state unfolding free energy of Δ*G*^*o*^ = 4.6 kcal mol^-1^ at 25 °C,^43^ and it is prone to resist aggregation; thus, it is well suited as a model protein for studies such as these. The strong curvature of the 50 nm nanoparticles is likely sufficient to destabilize the protein. Geometrically, this increase in size is significant. For example, if GB3 remains folded (with a hydrodynamic diameter of approximately 2.2 nm),^43^ approximately 11 layers of GB3 would be needed to account for Δ*d*_*H*_ on 50 nm PSNPs, and 1-2 layers would be needed on 200 nm PSNPs. Alternatively, the protein may be unfolded; in this case, a larger Δ*d*_*H*_ would require fewer GB3 molecules, since unfolded proteins occupy more hydrodynamic volume.^49^ Multiple layers of unfolded proteins could also be present.^50^ The dramatic difference between 50 nm and 200 nm suggests that the protein destabilization that occurs from adsorption is variable over this range of curvatures. This is interesting, because even at the size of 50 nm, the nanoparticle surface is effectively flat compared to the size scale of GB3.

An increase in nanoparticle size is also observed for the polystyrene-binding domain R2ab. Here, however, agglomeration is observed for all three sizes of nanoparticles. The observed polydispersity values from DLS are very high, and the hydrodynamic diameters in these experiments are not trustworthy; they are reported merely to demonstrate the substantial contrast to what is observed for GB3. We observed similar behavior for R2ab previously when working with 50 nm, carboxylate-functionalized PSNPs^36^; in that study, R2ab substantially increased ζ even at very low concentrations, reflecting a decreased colloidal stability of the PSNPs. R2ab is a basic protein, with a theoretical pI of 9.72 and a net charge of +11 at pH 7, and it can efficiently neutralize the surface charge on PSNPs. While the hydrodynamic radii are impossible to interpret quantitatively for R2ab, they nevertheless reflect a strong interaction between this protein and polystyrene; moreover, the interaction likely results in complex protein interactions as proteins are restructured on the nanoparticle surface.

### ITC Reveals a Significant Exothermic Binding for both GB3 and R2ab

The pattern observed for GB3 was intriguing, and we sought to probe the energetics and stoichiometry of these trends more deeply. To do this, we used isothermal titration calorimetry (ITC) which can provide reliable estimates of binding affinities and enthalpies between proteins and nanoparticles.^34,51,52^ By carefully controlling the concentration of proteins, we were able to avoid aggregation and obtain reproducible thermograms for both GB3 and R2ab with 50 and 100 nm PSNPs (**Fig. 1A-B** and Supporting Information, **Fig. S2**). We also extensively dialyzed all samples to eliminate the contribution from surfactants,^38^ and we confirmed that dilution heats for proteins and nanoparticles were negligible (Supporting Information, **Fig. S3**). For 200 nm PSNPs, severe aggregation was observed even at very low concentrations of R2ab (< 15μM), and only GB3 could be measured for this size.

Most of the adsorption profiles could be interpreted using a model with a single set of *N* identical, independent binding sites (**Fig. 1** and Supporting Information, **Table S4**). Clearly, the PSNP surface is complex and changes as the binding saturation increases; this model is a simplification. Nevertheless, such a model apparently describes the majority of interactions probed in this study. However, when GB3 was added to 50 nm PSNPs, a small endothermic hump was observed near the end of the titration (**Fig. 1A**). This hump was persistent and systematic over multiple protein preps, PSNP batches, and even ITC injection times. For the system in **Fig. 1A**, good agreement was obtained by fitting a model with two sets of *N* identical binding sites to this data (Δ_*bind*_*H*_1_, Δ_*bind*_*H*_2_, *K*_1_, *K*_2_, and *N*_1_, *N*_2_). For GB3 binding to 50 nm PSNPs, we therefore refer to two binding processes, a dominant exothermic process (Δ_*bind*_*H*_1_, *K*_1_, and *N*_1_) that occurs early in the titration, and an endothermic process that is not observed until the titration is nearly complete (Δ_*bind*_*H*_2_, *K*_2_, and *N*_2_; Supporting Information, **Table S5**). For all other PSNP systems, only a single exothermic process was observed, and these exothermic binding processes are used for comparison with the exothermic process of GB3 binding to 50 nm PSNPs. It is unclear what the endothermic process is, but it may correspond to multiple layers of binding or the changing nature of the binding as the nanoparticle approaches saturation. Alternatively, Velazquez-Campoy *et a*l. suggest endothermic humps like these may be due to cooperativity of binding.^53^ To our knowledge, the origin of this endothermic peak remains unexplained.

The exothermic binding processes of both GB3 and R2ab share similar features. Exothermic interactions are largely controlled by electrostatic, van der Waals, and hydrogen bonding interactions where ΔH < 0.^54^ Many recent studies have found that protein-nanoparticle binding is exothermic, although there are a few exceptions.^54^ While there is unlikely to be a well-defined binding site on the protein surface, the exothermic interaction may reflect an average orientation of approach.^44^ For both GB3 and R2ab, we observed that the magnitude of the enthalpy change decreased as a function of curvature (**Fig. 1D**). Similar to what was observed by DLS, the binding interactions appeared to become weaker for the larger nanoparticles. This observation suggests that surface curvature plays a significant role in influencing the strength of the protein-nanoparticle interaction. Furthermore, we observed that R2ab consistently exhibited a more exothermic enthalpy of binding compared to GB3. This more negative enthalpy of binding observed for R2ab suggests that this protein has a larger number of favorable surface interactions per protein, reflecting its larger size (17 kDa vs. 6 kDa).

### As Nanoparticle Curvature Decreases, the Apparent Binding Affinity Decreases, Reflecting Complex Changes in Enthalpy and Entropy

The observed enthalpies suggest that the strength of the protein-nanoparticle interaction becomes weaker as the curvature becomes less severe. This is also borne out by the apparent binding affinities for R2ab and GB3. In ITC, the measured binding affinity (*K*_*bind*_ or Δ_*bind*_*G*^*o*^) is an apparent affinity because it reflects the overall heat evolved during the binding process, including other processes such as conformational changes or competing interactions. While apparent binding affinities may reflect multiple processes, it is possible to compare these affinities provided the other factors do not change between experiments.^55,56^ This is expected to be the case for well-controlled titrations of proteins into nanoparticles. Analyzing the ITC data, we observed that larger-sized nanoparticles exhibit a decreased free energy of binding for both GB3 and R2ab **(Fig. 1C)**. These proteins bind more favorably to nanoparticles with greater surface curvature, and it is consistent with the decrease in the magnitude of the enthalpy observed as a function of nanoparticle size. Interestingly, the changes in Δ_*bind*_*G*^*o*^ do not strictly follow changes in Δ_*bind*_*H*; instead, the decrease in binding affinity includes contributions from both changing enthalpy and entropy contributions (**Fig. 1D-E**). In other words, both entropic and enthalpic contributions to binding become less favorable as the PSNP size increases, and protein disorder may play a role. It is also interesting to note that, while GB3 starts out as the more favorable of the two proteins for 50 nm PSNPs, its Δ_*bind*_*G*^*o*^ becomes less favorable relative to R2ab as a function of curvature; for 100 nm PSNPs, R2ab is the more favorable protein. The fact that R2ab retains high binding affinity as curvature decreases may explain its propensity to interact with flat polystyrene surfaces, driving biofilm formation. One might expect a simple adsorption process to be mainly enthalpic, with little change as the curvature is altered; here, these results implicate protein restructuring as well as simple binding.

### GB3 and R2ab are Well-Solvated, Disordered Polymers when Adsorbed to PSNPs

The thermodynamic parameters suggest that protein binding may induce unfolding in the PSNP corona; therefore, we decided to revisit our interpretation of the DLS data in light of the ITC-determined stoichiometries. Previously, we observed that the binding stoichiometry and Δ*d*_ℎ_ on gold nanoparticles could be predicted using a simple geometric model.^4,43^ In that model, the surface area occluded by a globular protein was used to predict the number of proteins that could fit on the nanoparticle surface, assuming monolayer coverage. Such a relationship does not appear to hold for PSNP binding. While the number of bound proteins (*N*) does increase as the nanoparticle size increases (Supporting Information, **Table S6**), in all cases, *N* is far less than what would be expected from geometrical considerations using the folded structure or even a hydrodynamic volume estimated from an unfolded polymer’s radius of gyration (**Fig. 1F**).^49^ This is particularly striking in light of the hydrodynamic diameters, which suggest that multiple layers of proteins are bound to the surface. While establishing the concentration of PSNPs is not as straightforward as it is for AuNPs,^57^ our concentrations are based on the manufacturer’s particle counts and gravimetric analysis; it is unlikely that the discrepancy originates from particle concentration. Instead, this discrepancy may be related to the loss of protein stability: both GB3 and R2ab are likely unfolded on the nanoparticle surface, sampling a disordered, but hydrated state. This state will exhibit a larger hydrodynamic volume than a compact, globular protein, or an unfolded protein that is flattened on the surface. The layer of disordered protein closest to the nanoparticle surface may then recruit other proteins, which can similarly unfold and adsorb to the growing protein-nanoparticle conjugate. This explanation fits both the large values observed for Δ*d*_*H*_ as well as the comparatively small values of *N*.

This model is also supported in part by the fact that the *N* values for GB3 are higher on each nanoparticle compared to R2ab by a factor of 3-4 (*N*_*GB*3_/*N*_*R*2*ab*_). The dimensions of unfolded proteins scale according to 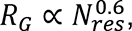 where *R*_G_ is the radius of gyration and *N*_*res*_ is the number of residues (**Fig. 1F**).^49,58^ The R2ab domain has 156 residues and GB3 has 56, and the ratio of their predicted radii is 1.85. Using these radii, one can estimate the occluded surface area on a nanoparticle (*A* = *πr*^2^). The number of adsorbed proteins is inversely proportional to this area; thus, for a well-solvated, disordered conformation, should be *N*_*GB*3_/*N*_*R*2*ab*_ ≈ 3.42. For globular proteins, 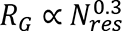,^59^ which leads to a predicted stoichiometric ratio of *N*_*GB*3_/*N*_*R*2*ab*_ ≈ 1.85. Thus, even though the observed stoichiometric cannot be predicted directly based on geometric considerations, the ratio *N*_*GB*3_/*N*_*R*2*ab*_ is more consistent with a hydrated, disordered polymer than a compact, globular protein. Moreover, a disordered polymer can more easily explain the large Δ*d*_*H*_ observed by DLS. Predicting the occluded surface area accurately is difficult because many factors can contribute to the hydrodynamic packing of disordered polymers. Comparing ratios removes some of these complications, such as the contribution from the persistence length,^60,61^ and this simple ratiometric approach also seems to support an extended, disordered protein structure in the PSNP corona.

### PSNPs Destabilize the Folded State of R2ab and GB3, with Higher Curvature Destabilizing Proteins to a Greater Extent

The DLS and ITC data suggest that adsorption to PSNPs favors an unfolded state for GB3 and R2ab. We sought to test this directly by measuring protein stability in the presence of PSNPs. As with the DLS experiments, the total PSNP surface area was held constant for all three sizes of PSNP, and the nanoparticle concentration was set low enough to minimize aggregation while ensuring that a significant population of protein was interacting with the PSNP surface. Protein stability was measured using intrinsic tryptophan fluorescence and increasing the guanidinium chloride (GdmCl) concentration (**Fig. 2A**), and the linear extrapolation model was used to extract Δ_*unfold*_*G*^*o*^(*H*_2_*O*) and an *m*-value.^42^ This approach has been employed previously with Ribonuclease A on silica nanoparticles.^62^ In that work, the authors found that ultrasmall (4 nm) particles perturbed the unfolding free energy less than larger (15 nm) nanoparticles. Here, we examine nanoparticles that are much larger than the size of the proteins themselves (50-200 nm).

**Figure 2.**
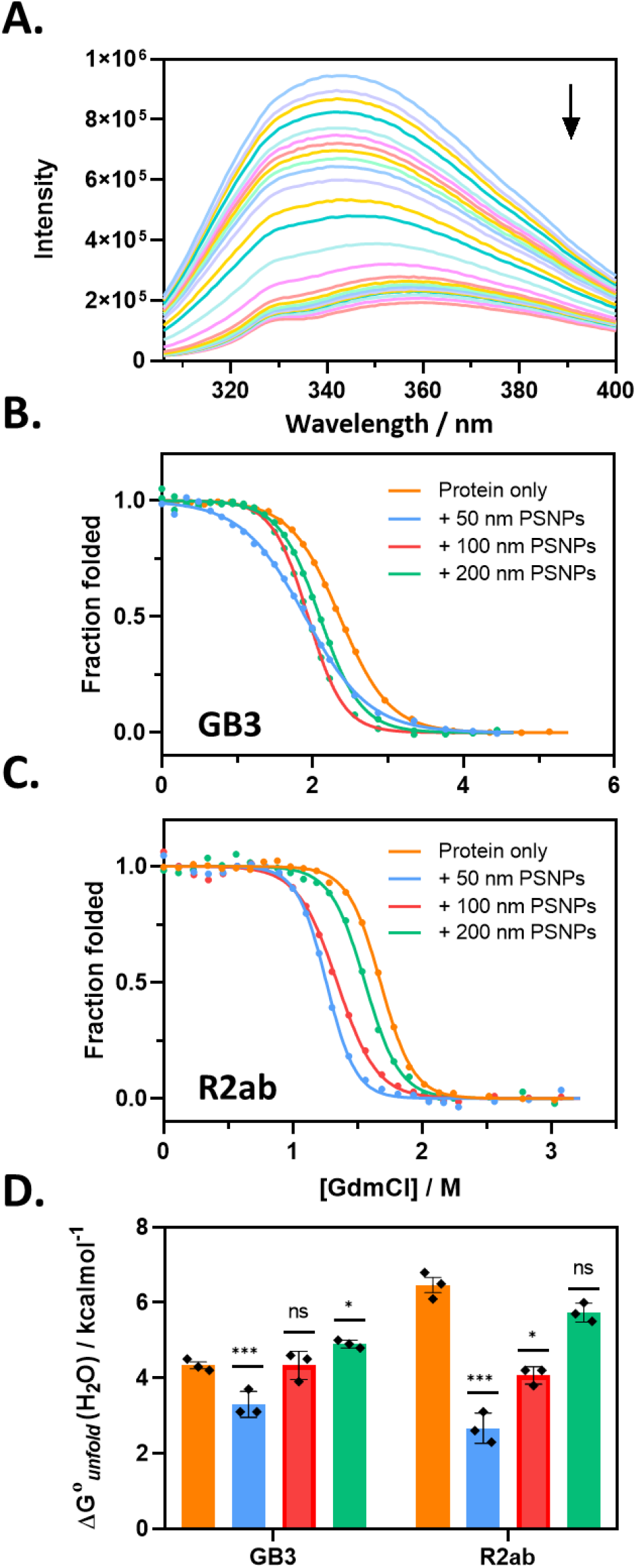
Protein stability in the PSNP corona is perturbed most for smaller nanoparticles. (A) shows a typical Trp fluorescence spectrum as GdmCl is added for R2ab. The arrow indicates increasing GdmCl. Fluorescence denaturation monitored at 344 nm following baseline correction for GB3 (B) and R2ab (C). For both proteins, the stability is most perturbed in the presence of 50 nm PSNPs. Summary of Δ_*unfold*_*G*^*o*^ for each combination of protein and nanoparticle is shown in (D). The statistical significance of the difference relative to the GB3 or R2ab in the absence of PSNPs is shown. Significance relative to the no-nanoparticle value was assessed using one-way ANOVA with Tukey’s multiple comparisons test (n.s., not significant; *, p < 0.05, ***, p < 0.001. Error bars represent the standard error of the mean of three experiments. For (B-D), colors represent no nanoparticles (orange), 50 nm PSNPs (blue), 100 nm PSNPs (red), 200 nm PSNPs (green).

For both GB3 and R2ab on 50 nm PSNPs, we observe a significant decrease in Δ_*unfold*_*G*^*o*^(*H*_2_*O*), similar to what was observed for 15 nm silica nanoparticles. However, as the curvature decreases, we observe that the net destabilization also decreases (**Fig. 2B-D** and Supporting Information, **Table S7**) and in the presence of 200 nm PSNPs, the stabilities for GB3 and R2ab are statistically identical to their stabilities in the absence of nanoparticles. Together with the RNase A work, this suggests that small nanoparticles, similar in size to proteins themselves, may not affect protein stability significantly, but stability approaches a minimum for larger, curved nanoparticles. As nanoparticles continue to increase in size, the nanoparticle appears more like a flat surface, and the impact on stability once again becomes marginal.

The observed values of Δ_*unfold*_*G*^*o*^ are in good agreement with expectations from binding experiments and DLS data. The surfaces with the greatest curvature in our study are those with the highest value of Δ_*bind*_*G*^*o*^, and large binding free energies can drive protein unfolding. Correspondingly, large perturbations to Δ_*unfold*_*G*^*o*^ will be associated with changes in average ensemble dimensions, and these changes will influence the binding stoichiometry and the hydrodynamic diameter. The linear extrapolation model also produces *m*-values as part of the fitting process; these *m*-values roughly correlate with the size of the domain involved in the folding transition.^63^ The *m*-values we observe (Supporting Information, **Table S7**) follow a similar trend to Δ_*unfold*_*G*^*o*^, although the statistical differences are small.

### When Total Nanoparticle Surface Area Is Held Constant, Smaller, High-Curvature PSNPs Disrupt the Protein Structure More Than Larger PSNPs

Our binding and folding stability measurements suggest that PSNPs can unfold proteins, with smaller nanoparticles having a more significant effect. To confirm this, we used far-UV circular dichroism (CD) to monitor protein secondary structure.^64,65^ A 0.2 mm pathlength demountable cuvette was used to reduce the influence of scattering for larger particles. Overall, the CD spectra are consistent with our observations above: When controlled for total surface area, 50 nm PSNPs were more disruptive to protein secondary structure than 100 and 200 nm PSNPs (**Fig. 4A-B**). For GB3, the molar ellipticity was severely affected for 50 and 100 nm PSNPs; however, when controlling for the total PSNP surface area, the spectrum in the presence of 200 nm PSNPs was very close to the spectrum in the absence of nanoparticles. The same trend holds for R2ab; however, the 50 nm PSNPs appear to be less disruptive to the secondary structure for this protein, suggesting that a larger population of R2ab may retain its folded structure, even for the smallest PSNPs tested. This feature may explain why R2ab is well suited for binding polystyrene surfaces.^36^ It is impossible to interpret secondary structures quantitatively in exchanging populations of proteins;^64^ while populations of helix, strand, and coil are frequently reported for protein-nanoparticle mixtures, these are almost assuredly inaccurate when exchange with the surface is present. Nevertheless, it is clear that the native structure is progressively less affected for GB3 and R2ab as the curvature decreases. Loss of secondary structure has been observed for many protein corona systems prior to this,^66,67^ but here we have shown that structural disruption tracks closely with energetic trends as a function of curvature.

To further explore the structural perturbations in GB3 and R2ab, we monitored hydrophobic group exposure with 8-anilino-1-naphthalenesulfonic acid (ANS), a fluorescent solvatochromatic dye. The ANS fluorescence profile and its peak maximum changes upon the dye’s non-covalent interaction with protein hydrophobic regions.^68^ Generally, ANS becomes more fluorescent as a protein unfolds and hydrophobic surfaces are exposed. By adding ANS to protein-PSNP solutions, we were able to monitor the effect of nanoparticle size on hydrophobic surface exposure (**Fig. 3C-D**). In agreement with CD, we found that the nanoparticle size had a significant effect on the ANS fluorescence for both proteins analyzed. When corrected for intrinsic ANS-PSNP fluorescence, both R2ab and GB3 exhibited similar trends for ANS fluorescence intensities: the overall ANS intensity was reduced as the size of the nanoparticles increased. For both proteins, in the presence of 50 nm PSNPs, the highest fluorescence intensity was observed, corresponding to maximal hydrophobic exposure and consistent with an extended, solvated protein adsorbed to the PSNP surface.

**Figure 3.**
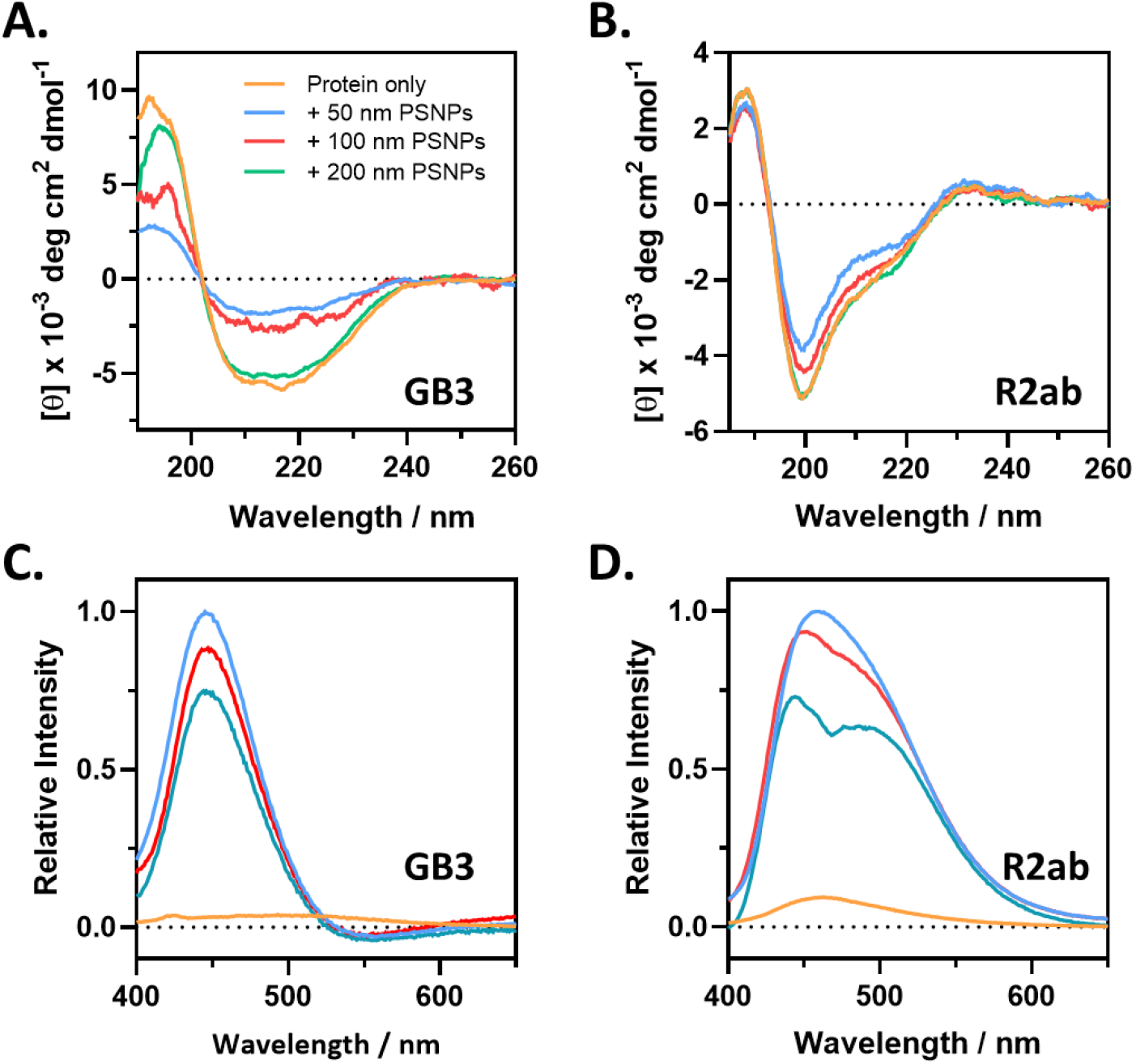
When controlled for total PSNP surface area, structural perturbations in the corona are most pronounced for smaller nanoparticles. GB3 (A) and R2ab (B) exhibit characteristic CD spectra in the presence of increasing sizes of nanoparticles. In the absence of nanoparticles (orange), both proteins exhibit a well-defined secondary structure. Normalized ANS fluorescence spectra for GB3 (C) and R2ab (D), in the presence of 50 nm (blue), 100 nm (red) and 200 nm (green) PSNPs show emission in the range 400 to 650 nm. In the absence of nanoparticles, both proteins exhibit minimal ANS fluorescence in their folded state, since hydrophobic patches are buried. Hydrophobic groups are exposed for 50 nm PSNPs, but as nanoparticle size is increased, the fluorescence intensity is reduced due to reduced exposure of hydrophobic groups. Note that the negative fluorescence in panel (C) arises from imperfect cancellation of the experimental spectrum and the ANS-PSNP control (Supporting Information, **Fig. S4**).

Protein rearrangements and conformational changes on a nanoparticle are common phenomena in the protein adsorption process.^69–71^ Here, the ANS approach both supports and complements the observations from CD spectroscopy. First, the trends of decreasing structural perturbation with increasing nanoparticle size is consistent for both proteins. Second, even though the CD spectrum of R2ab is similar to the native spectrum in the presence of 200 nm PSNPs, there is still apparently significant hydrophobic exposure. This may reflect a “molten-globule” like structure or a conformation where the two subdomains of R2ab are decoupled. The presence of a shoulder at 460 nm is also evident for R2ab on 200 nm PSNPs; this effect for ANS has been observed previously^72,73^ and could indicate a conformational change. It is challenging to interpret these features quantitatively because the hydrophobic patches exposed to solvent are not linearly correlated to the fraction of protein unfolded. Nevertheless, it is clear that CD spectroscopy and ANS fluorescence exhibit similar trends with respect to nanoparticle curvature. Moreover, even when the CD spectrum in the presence of nanoparticles is similar to the native CD spectrum, there may still be structural perturbations present in the adsorbed population.

### Trends in Protein Structural Changes Can Be Extrapolated to Macroscopic Surfaces on Treated Polystyrene Plates

Finally, we sought to investigate whether the observations made for PSNPs could be extrapolated to cell culture-treated polystyrene plates. There are many challenges to studying protein behavior on macroscopic surfaces. First, there are few biophysical tools to study the structure and properties of proteins adsorbed to a macroscopic surface. Second, macroscopic surfaces have a low surface to volume ratio, and this affects sensitivity, making detection of the adsorbed protein difficult. Moreover, comparisons between nanoparticles and macroscopic surfaces can be complicated by differing surface chemistries,^74^ variations in surface nanostructure,^27^ and the presence of surfactants used to stabilize colloidal particles.^38^ Nevertheless, it is possible some similarities exist as well, especially as curvature approaches a limiting value. Having investigated GB3 and R2ab binding to PSNPs, we sought to compare how these proteins bind to treated polystyrene cell culture plates. These plates were chosen because of their similar chemistry and their comparable zeta potential relative to PSNPs,^75^ and their ubiquitousness in biomedical diagnostics.

We noticed during our experiments with ANS that, even in the absence of PSNPs, some fluorescence was detectable when adding protein to polystyrene plates. No fluorescence was observable in quartz cuvettes or polypropylene centrifuge tubes. This suggested that ANS could be used to monitor structural perturbations of GB3 and R2ab adsorbed on macroscopic polystyrene surfaces. Using ANS and buffer (no protein) as a reference, we determined the observable fluorescence at 480 nm after one hour of incubation for protein in a 6-well culture plate (**Fig. 4A** and **Fig. 4B-C**, “Plate Only”). Then, solutions were prepared in similar plates with PSNPs, such that the total PSNP surface area was approximately equal to the surface area of a single well in a 6-well plate (see Methods for more details). For these solutions, a reference containing PSNPs, ANS, and buffer was used. Once again, the fluorescence at 480 nm was detected after 1 hr incubation with PSNPs and protein (**Fig. 4B-C**, “50-200 nm PSNPs”). These experiments enable comparing adsorption to the plate alone relative to the adsorption of a similar amount of surface on PSNPs. There are several challenges in interpreting this comparison quantitatively. First, even though we have controlled for the total surface area (plate vs. PSNPs), the polystyrene plate is not perfectly smooth, and the comparison is only approximate. Second, as stated before, ANS fluorescence is not linearly proportional to protein disorder. Nevertheless, once again, a trend is observed. More ANS fluorescence is present for smaller, high curvature nanoparticles, and a smaller but detectable ANS fluorescence is present for the plates alone. The trend in ANS fluorescence follows the trend we observed for nanoparticles alone in quartz cuvettes: adsorption on flatter surfaces appears to be less disruptive to protein structure than adsorption to highly curved surfaces. Alternatively, the binding affinity may be lower for flat surfaces, which would also corroborate the trends in our biophysical data. In this case, the lower ANS fluorescence would reflect a lower population of surface-bound protein.

**Figure 4.**
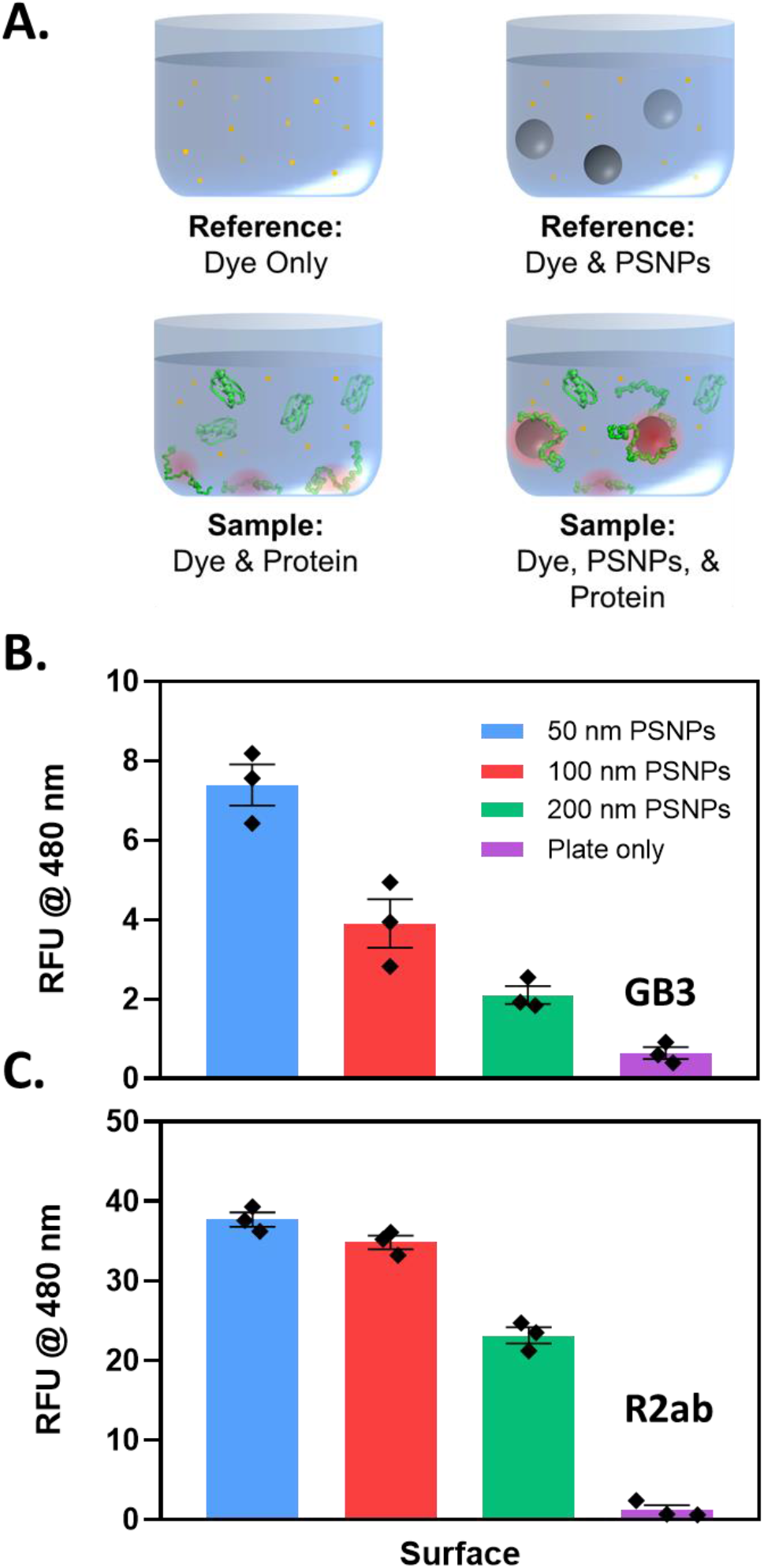
A small but detectable ANS fluorescence signal is present even on flat polystyrene plates, consistent with trends for PSNP curvature. (A) Subtracting out a reference with no protein (top left) allows visualization of fluorescence with protein adsorbed to a cell culture plate surface (bottom left). Similarly, subtracting out a reference with PSNPs and no protein (top right) enables fluorescence to be measured when protein adsorption is present (bottom right). The total PSNP concentration is set so that the PSNP surface area and culture plate surface are comparable. The behavior of GB3 (B) and R2ab (C) follow similar trends, where ANS fluorescence is measured as relative fluorescence units (RFU) at a λ_max_ of 480 nm. The lowest ANS fluorescence is observed for flat polystyrene surfaces, whereas fluorescence increases significantly for curved nanoparticles.

## Discussion

Protein structure in a nanoparticle corona will determine which bio-nano interactions are important for that nanoparticle. If proteins remain globular in the corona, their orientations and packing will establish the set of surfaces that can be detected by the immune system. On the other hand, if proteins in the corona are disordered, the orientation is ill-defined; instead, protein flexibility and hydration will determine the relevant interactions. Here, we see evidence for a disordered, well-hydrated corona on PSNPs, as opposed to a densely packed multilayer of unfolded proteins. Our model is supported by a relatively low protein-nanoparticle binding stoichiometry, a large increase in nanoparticle hydrodynamic diameter in the presence of protein, and a marked destabilization of proteins in the presence of PSNPs. Significant structural perturbation (as monitored by CD) and hydrophobic exposure (as monitored by ANS fluorescence) indicate that many residues are solvent accessible. At the same time, the strong enthalpy of interaction suggests that proteins in the corona are anchored tightly to the nanoparticle surface. Together, these data paint a picture where loops of flexible, disordered protein are separated by regions that interact with the PSNP surface (**Fig. 5**). We propose that these anchor points help to stabilize the corona, interacting with the nanoparticle surface much like epitopes bind to an antibody. While the interaction is likely weaker and more fluid, these hypothesized “adsorbotope” sequences are ultimately responsible for the association between the nanoparticle and the protein. In this model, the exposed loops are able to interact with other proteins, including opsonins and other immune system components, ultimately determining the nanoparticle fate. For larger PSNPs, the protein remains disordered, as evidenced by ANS fluorescence, but the structure may be more similar to a molten globule, with less exposed surface.^76^ For smaller PSNPs, more disorder is present, and this may lead to additional multilayer binding events such as those observed for GB3 on 50 nm PSNPs.

**Figure 5.**
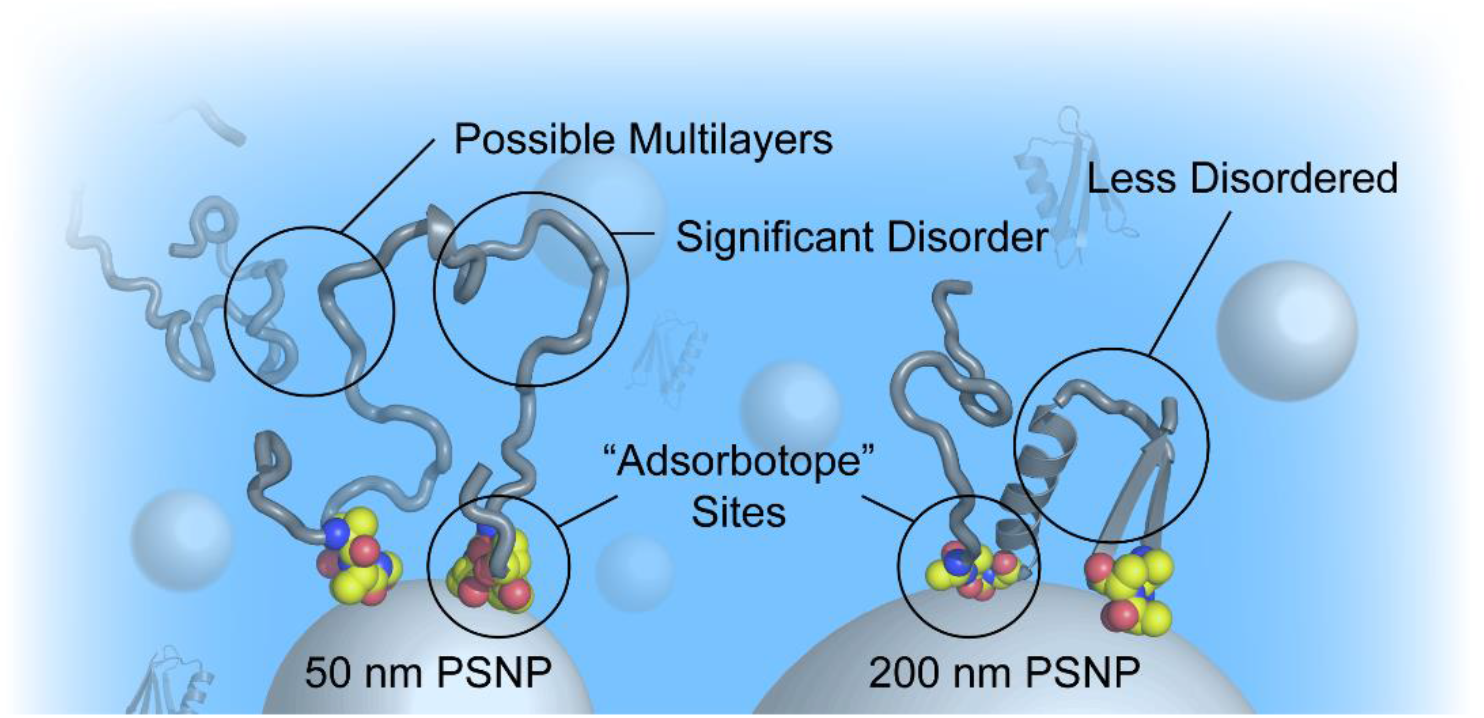
A model for disordered coronas on PSNPs. Regions of the protein that favorably interact with the nanoparticle surface are shown as CPK spheres (carbon is colored yellow). Other well-solvated, disordered protein regions are represented as grey cartoons. The degree of disorder decreases as the nanoparticle curvature is reduced; at high curvatures, the disorder my promote multilayer interactions.

The possibility of distinct protein regions that favor binding to a nanoparticle surface is supported by several recent studies of protein and peptide interactions with nanoparticles. For example, NMR saturation transfer experiments by Zhang, *et al.* demonstrate that certain amino acid side chains, such as Trp and Phe, exhibit strong π-π interactions with a polystyrene surface.^77^ In a follow-up study, Xu, *et al.* found that non-natural amino acids with long aliphatic chains also favored binding to PSNPs.^78^ The implication of these investigations is that sequences containing a high fraction of hydrophobic and aromatic side chains could form the basis of adsorbotope sites. Indeed, regions of high affinity were observed for a p53 fragment and α-synuclein in the presence of silica nanoparticles, and a free-residue interaction model has been developed to characterize sequence binding specificity.^79,80^ The p53 and α-synuclein proteins are already disordered, but a similar approach could be used to identify potential adsorbotopes on PSNPs. Presumably, identifying nanoparticle-binding regions could enable the prediction of unbound protein loops that can interact with cell receptors and immune system components. Such an approach could be used to predict the transport mechanisms by which nanoparticles are internalized in cells.^81,82^

Finally, it has been observed that nanoparticles can modulate protein aggregation or the formation of amyloid fibrils.^83–86^ Two studies stand out for PSNPs, and our model conveniently explains the observed behavior. In the first study, amine-functionalized PSNPs were shown to both accelerate and slow fibrillation of the amyloid β (1-40, Aβ) peptide depending on the protein:surface ratio.^87^ At a fixed surface area of 1 m^2^ L^-1^ and similar protein concentrations (8 μM), larger PSNPs (120-140 nm) increased Aβ fibrillation rates relative to smaller PSNPs (57 nm). Thus, under similar conditions, larger PSNPs favored faster fibrillation. This is readily explained by reduced protein destabilization on larger PSNPs, as observed in our experiments (**Fig. 2**): the highly destabilized proteins on smaller PSNPs are more random and are less competent to nucleate fibrils. A second study demonstrated that larger PSNPs (400 nm) were able to more readily induce fibrillation in hen egg white lysozyme compared to smaller PSNPs (100 nm).^88^ The authors of this study hypothesized that structural differences in adsorbed proteins can lead to different fibrillation rates. Our work suggests that, while lysozyme is destabilized overall by PSNPs, it is slightly more stable on larger particles relative to smaller particles. On larger particles, lysozyme can more effectively nucleate fibrils. The specific fibrillation behavior will depend on both the protein identity and the nanoparticle surface chemistry, but our model appears to be consistent with prior studies of fibrillation in the presence of PSNPs.

## Conclusion

In this study, we conducted an investigation to understand the impact of nanoparticle surface curvature on protein binding and corona interactions. Our focus was on R2ab, a bacterial surface protein derived from *S. epidermidis*, known for its strong affinity to polystyrene surfaces. To establish a baseline for comparison, we also studied GB3, a model protein. Surprisingly, we observed the same trend for both proteins: binding affinity decreased as the nanoparticle size increased, and, when controlled for total surface area, larger nanoparticles appeared to destabilize both proteins less than smaller particles. Given the large increase in hydrodynamic size relative to the number of proteins bound to each nanoparticle, we find evidence for a highly disordered, solvent accessible protein corona on PSNPs. Protein disorder is apparent for all sizes of PSNPs studied, but it appears to decrease on larger-sized PSNPs with less curvature. This finding has significant implications for protein accessibility and structure in coronas where disorder is implicated. Our data also provide insight into how proteins may interact with a flat macroscopic surface. Specifically, we hypothesize the existence of peptide segments with high nanoparticle binding affinity (“adsorbotopes”) interspersed with exposed, disordered protein segments. This model of protein structure on nanoparticle surfaces appears to rationalize fibrillation behavior in the presence of PSNPs. Moreover, it may help explain how cells interact with different-sized nanoparticles in biological systems.

## Associated Content

The Supporting Information is available free of charge at…

- Nanoparticle characterization (Tables S1-3, Figure S1)
- Tables of thermodynamic data (Table S4-5, Table S7) and representative ITC thermograms (Figures S2-3)
- Predicted adsorption stoichiometries assuming a globular protein (Table S6)
- Representative ANS fluorescence data (Figures S4-5)
- All raw experimental data has been uploaded to Zenodo (https://doi.org/10.5281/zenodo.8105819)

## Notes

The authors declare no competing financial interest.

## Acknowledgments

We thank Tim Dowell for a helpful presentation on crafting scientific illustrations. This work was supported by the National Institute of Allergy and Infectious Diseases of the National Institutes of Health under grant number R01AI139479 and the National Science Foundation under grant number MCB 1818090.

