## Supplemental Information for "Protein Binding Leads to Reduced Stability and Solvated Disorder in the Polystyrene Nanoparticle Corona"

### Supporting Tables

**Table S1. Protein and nanoparticle concentrations used in DLS and zeta potential measurements.**

| | Approximate Concentrations |
|---|---|
| R2ab (μM) | 15.0 |
| GB3 (μM) | 40.0 |
| 50 nm PSNPs (nM) | 53.0 |
| 100 nm PSNPs (nM) | 13.2 |
| 200 nm PSNPs (nM) | 3.3 |

**Table S2. Final PSNP concentrations used for ANS-plate experiments.**

Experiments used a final volume of 0.5 mL.

| PSNP size (nm) | Concentration (nM) |
|---|---|
| 50 | 0.40 |
| 100 | 0.10 |
| 200 | 0.025 |

**Table S3. Observed zeta potentials for PSNPs.**

| PSNP size (nm) | 50 | 100 | 200 |
|---|---|---|---|
| Zeta Potential (mV) | -48 ± 4 | -53 ± 4 | -27 ± 6 |

**Table S4. Thermodynamic parameters for interactions of GB3 and R2ab with PSNPs.**

| Parameter | GB3/50 nm | GB3/100 nm | GB3/200 nm | R2ab/50 nm | R2ab/100 nm |
|---|---|---|---|---|---|
| $K_{bind,app}$ | (5.2 ± 0.4) x $10^8$ | (1.9 ± 0.6) x $10^7$ | (1.9 ± 0.6) x $10^5$ | (1.5 ± 0.1) x $10^8$ | (7.2 ± 0.4) x $10^7$ |
| $N$ | 580 ± 2 | 1037 ± 19 | 1631 ± 12 | 197 ± 6 | 241 ± 8 |
| $\Delta_{bind}H$ (kcal $mol^{-1}$) | -24.6 ± 1.4 | -20.0 ± 1.3 | -9.4 ± 1.3 | -27.6 ± 1.1 | -22.2 ±1.1 |
| $\Delta_{bind}G^o$ (kcal $mol^{-1}$) | -11.8 ± 0.9 | -9.9 ± 0.4 | -7.7 ± 0.5 | -11.1 ± 0.6 | -10.6 ± 0.5 |
| $-T\Delta_{bind}S$ (kcal $mol^{-1}$) | 12.8 ± 1.4 | 10.1 ± 1.3 | 1.7 ± 1.3 | 16.5 ± 1.1 | 11.6 ± 1.1 |

**Table S5. Thermodynamic parameters for GB3 binding to 50 nm PSNPs (second site).**

| Parameter | Process 1 | Process 2 |
|---|---|---|
| $K_{bind,app}$ | (5.2 ± 0.4) x $10^8$ | (1.1 ± 0.2) x $10^7$ |
| $N$ | 580 ± 2 | 290 ± 2 |
| $\Delta_{bind}H$ (kcal mol$^{-1}$) | -24.6 ± 1 | 10.3 ± 3 |

**Table S6. Comparison of N values for proteins bound to nanoparticles.**

Predictions in the second column are adapted using the approach of Wang *et al.*[1]

| | N (ITC) | N (Wang et. al) |
|---|---|---|
| GB3/50 nm | 580 ± 2 | 2267 |
| GB3/100 nm | 1037 ± 19 | 9070 |
| GB3/200 nm | 1631 ± 12 | 36281 |
| R2ab/50 nm | 197 ± 6 | 797 |
| R2ab/100 nm | 241 ± 8 | 3191 |

**Table S7. Protein unfolding stabilities measured in the presence of PSNPs.**

| | **GB3** | | **R2ab** | |
|---|---|---|---|---|
| Sample | $\boldsymbol{\Delta G^{o}_{unfold}(H_2O)}$ (kJ mol$^{-1}$) | GdmCl $\boldsymbol{m}$ (kcal mol$^{-1}$ M$^{-1}$) | $\boldsymbol{\Delta G^{o}_{unfold}(H_2O)}$ (kJ mol$^{-1}$) | GdmCl $\boldsymbol{m}$ (kcal mol$^{-1}$ M$^{-1}$) |
| Protein only | 4.4 ± 0.1 | 2.1 ± 0.4 | 6.1 ± 0.8 | 4.9 ± 0.3 |
| + 50 nm PSNPs | 3.6 ± 0.5 | 1.8 ± 0.5 | 3.1 ± 0.6 | 2.2 ± 0.4 |
| + 100 nm PSNPs | 4.7 ± 0.5 | 2.2 ± 0.4 | 4.2 ± 0.2 | 2.7 ± 0.1 |
| + 200 nm PSNPs | 4.8 ± 0.8 | 2.8 ± 0.3 | 6.0 ± 0.8 | 4.4 ± 0.1 |

### Supporting Figures

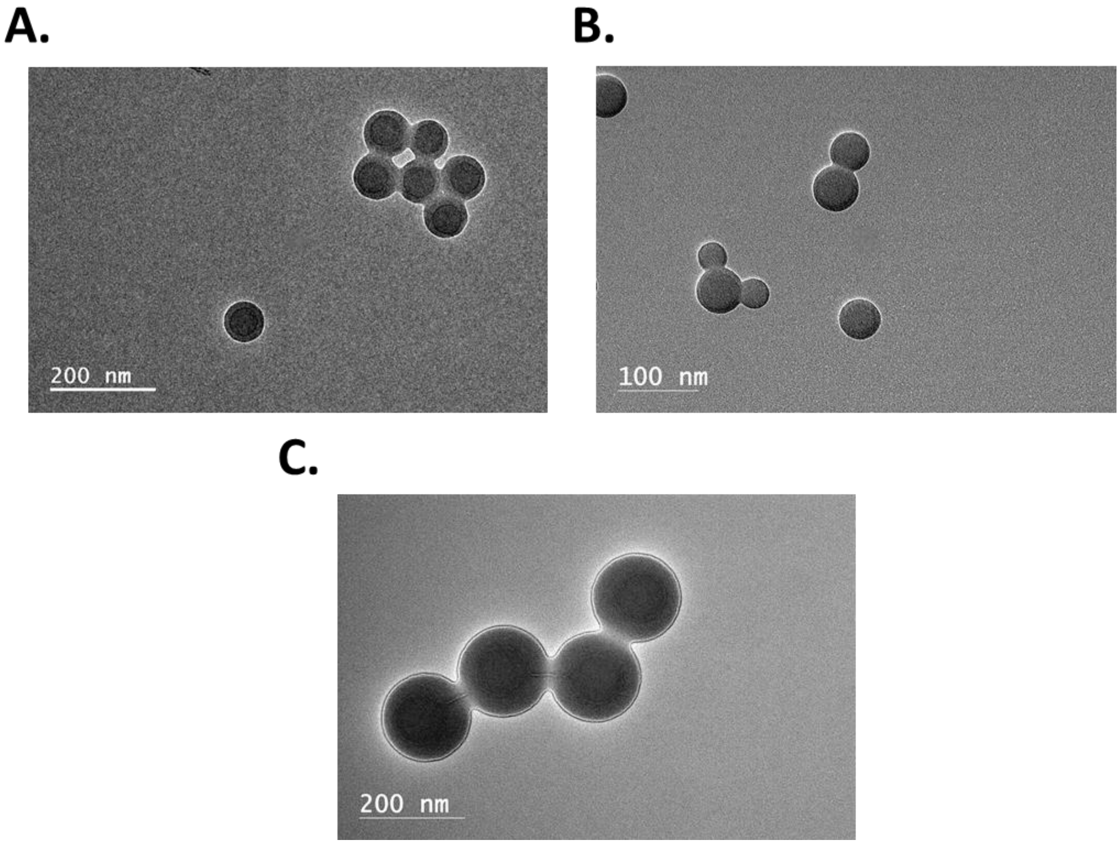

**Figure S1. TEM images representing non-functionalized polystyrene nanoparticles.**

Data is shown for 50 nm (A), 100 nm (B) and 200 nm (C)

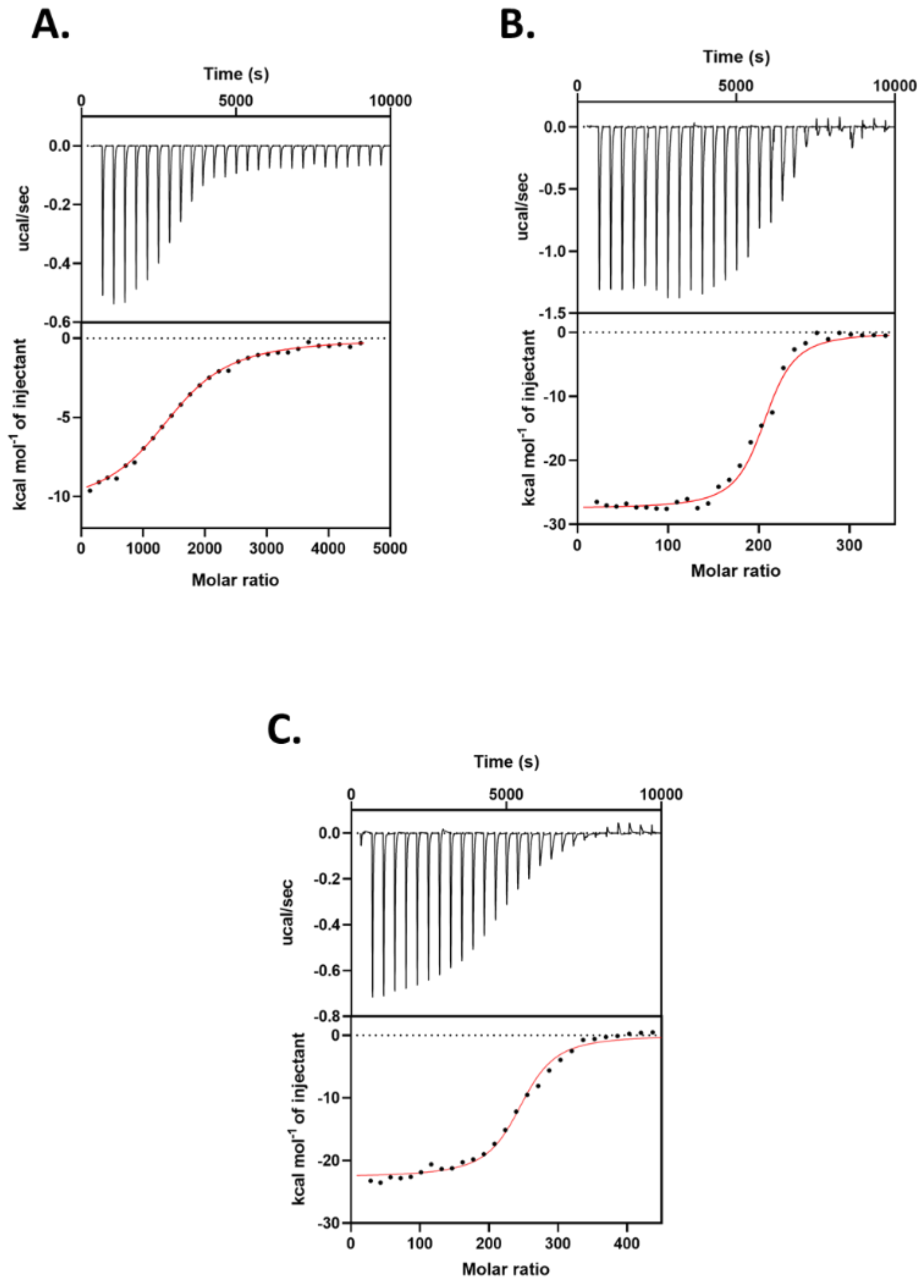

**Figure S2. Representative ITC profiles for proteins in the presence of PSNPs**

Data is shown for GB3/200 nm (A), R2ab/50 nm (B) and R2ab/100 nm (C)

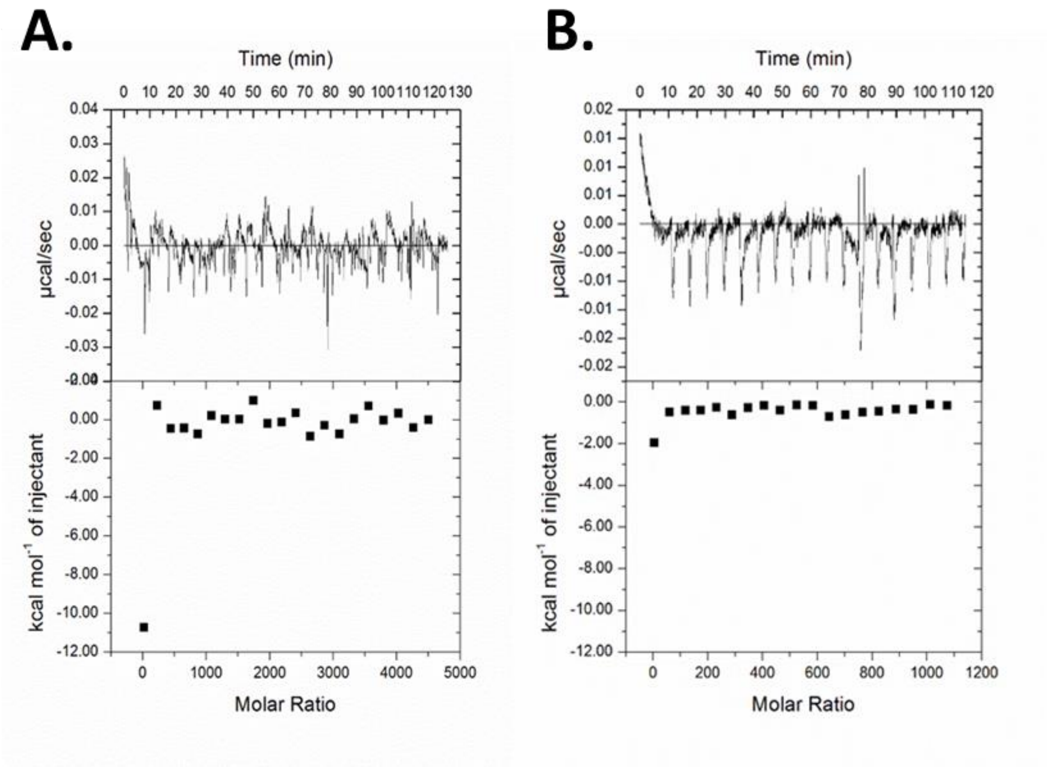

**Figure S3. Representative ITC heat profiles showing minimal heats of dilution.**

Dilution heats are shown for 50 nm PSNPs (A) and GB3 (B), upon the titration of the buffer.

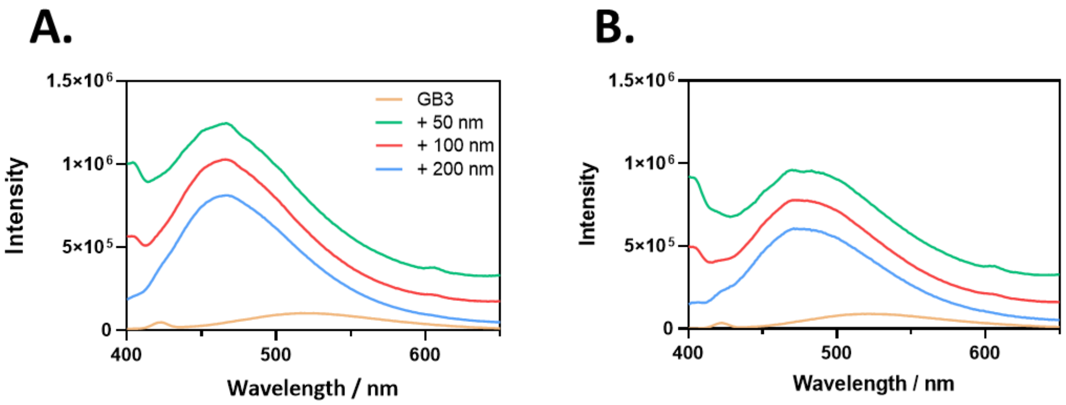

**Figure S4. ANS flouresence profiles for GB3 in the presence of PSNPs, prior to blank subtraction.**

GB3 in the presence of PSNPs and ANS (A) show the variation of fluorescence intensities based on the PSNP size, and ANS shows high affinity to PSNPs (B), thus serving as a blank.

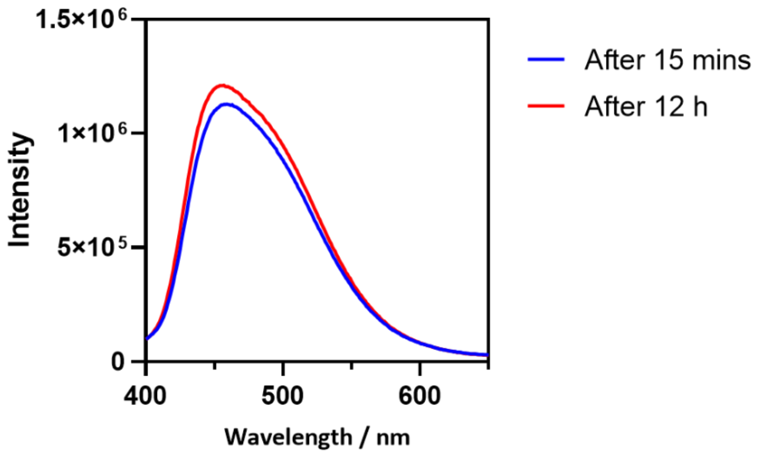

**Figure S5. Time dependence of ANS fluorescence profiles for R2ab in the presence of 50 nm PSNPs**

Spectra are shown after 15 minutes vs. after 12 hours of incubation in the dark. Both the spectra show no significant difference indicating that any exposure of hydrophobic patches due to unfolding has not progressed.

#### Supporting Information References

(1) Wang, A.; Vangala, K.; Vo, T.; Zhang, D.; Fitzkee, N. C. A Three-Step Model for Protein–Gold Nanoparticle Adsorption. *J. Phys. Chem. C* **2014**, *118* (15), 8134–8142. https://doi.org/10.1021/jp411543y.
